# Songbirds avoid the oxidative stress costs of high blood glucose levels

**DOI:** 10.1101/2023.05.09.539961

**Authors:** Csongor I. Vágási, Orsolya Vincze, Marie Adámková, Tereza Kauzálová, Ádám Z. Lendvai, Laura Pătra□, Janka Pénzes, Péter L. Pap, Tomáš Albrecht, Oldřich Tomášek

**Affiliations:** Evolutionary Ecology Group, Hungarian Department of Biology and Ecology, Babeş-Bolyai University, Cluj-Napoca, Romania; Department of Tisza Research, MTA Centre for Ecological Research-DRI, Debrecen, Hungary; Institute of Vertebrate Biology of the Czech Academy of Sciences, Brno, Czech Republic; Department of Botany and Zoology, Faculty of Science, Masaryk University, Brno, Czech Republic; Department of Zoology, Faculty of Science, Charles University, Prague 2, Czech Republic; Department of Evolutionary Zoology, University of Debrecen, Debrecen, Hungary; Department of Molecular Biology and Biotechnology, Centre of Systems Biology, Biodiversity and Bioresources, Babeş-Bolyai University, Cluj-Napoca, Romania

**Keywords:** Antioxidants, Glucose, Hyperglycaemia, Lipid peroxidation, Phylogenetic comparison, Physiological ecology

## Abstract

Chronically high blood glucose levels (hyperglycaemia) can compromise healthy ageing and lifespan at the individual level. Elevated oxidative stress can play a central role in hyperglycaemia-induced pathologies. Nevertheless, the lifespan of birds shows no species-level association with blood glucose. This suggests that the potential pathologies of high blood glucose levels can be avoided by adaptations in oxidative physiology at the macroevolutionary scale. However, this hypothesis remains unexplored. Here, we examined this hypothesis using comparative analyses controlled for phylogeny, allometry and fecundity based on data from 51 songbird species (681 individuals with blood glucose and 1021 individuals with oxidative state data). We measured blood glucose at baseline and after stress stimulus and computed glucose stress reactivity as the magnitude of change between the two time points. We also measured three parameters of non-enzymatic antioxidants (uric acid, total antioxidants and glutathione) and a marker of oxidative lipid damage (malondialdehyde). We found no clear evidence for blood glucose concentration being correlated with either antioxidant or lipid damage levels at the macroevolutionary scale, as opposed to the hypothesis postulating that high blood glucose levels entail oxidative costs. The only exception was the moderate evidence for species with a stronger stress-induced increase in blood glucose concentration evolving moderately lower investment into antioxidant defence (uric acid and glutathione). Neither baseline nor stress-induced glucose levels were associated with oxidative physiology. Our findings support the hypothesis that birds evolved adaptations preventing the (glyc)oxidative costs of high blood glucose observed at the within-species level. Such adaptations may explain the decoupled evolution of glycaemia and lifespan in birds and possibly the paradoxical combination of long lifespan and high blood glucose levels relative to mammals.

**Summary statement:** High blood glucose levels can harm organisms by causing oxidative stress. We show that, at the macroevolutionary level, songbirds defy this expectation, as their glucose levels and oxidative physiology are uncoupled.

## INTRODUCTION

Living organisms use chemical energy to sustain their vital functions, with carbohydrates, lipids, and proteins as the main metabolic energy substrates. Among carbohydrates, glucose has a prominent role in metabolism as it fulfils multiple metabolic functions. Besides being an important energy carrier, it is a vital precursor in the synthesis of a wide variety of other important biomolecules, including fatty, amino, and nucleic acids, cholesterol, glycoproteins, and glycolipids (Braun and Sweazea, 2008; Sweazea, 2022). The mere existence of several regulatory mechanisms that control glucose metabolism both at baseline conditions and during perturbations (e.g. stress response) provides evidence for the biological significance of glucose homeostasis (Braun and Sweazea, 2008). Deregulation of glucose homeostasis induces metabolic stress with a plethora of deleterious health consequences (Picard et al., 2014).

Birds have outstandingly high blood glucose levels in normal physiological state, in average two-to four times higher than that of similar-sized mammals (Braun and Sweazea, 2008; Holmes et al., 2001; Satoh, 2021). Only frugivorous and nectarivorous bats with a sugar-rich diet can have blood glucose values comparable to some birds (Kelm et al., 2011; Peng et al., 2017). Constitutively high blood glucose in birds is evolutionarily coupled with athletic capacity, high metabolic rate and elevated body temperature (Satoh, 2021). Moreover, avian blood glucose levels coevolve positively with fecundity, suggesting its importance in sustaining intense reproduction (Tomášek et al., 2019; Tomášek et al., 2022). Directional selection therefore might drive the evolution of high blood glucose until it incurs physiological costs that set an upper limit for circulating glucose levels.

Constitutively high blood glucose levels in birds are coupled, at least partly, with their evolved insulin resistance and lower insulin/insulin-like growth factor 1 signalling (Satoh, 2021). Remarkably, the blood glucose levels at physiological baseline of birds would be considered hyperglycaemia in humans and laboratory mammals, where chronic hyperglycaemia is a major health complication of diabetes and is causally involved in a variety of pathologies associated with this disease (Sweazea, 2022). Oxidative stress is believed to be the unifying mechanism behind most of the deleterious consequences of high blood glucose levels (Brownlee, 2001). Oxidative stress is a condition when an excessive production of pro-oxidants (i.e. beyond the low amounts needed for the body’s biological functions, such as gene translation or white blood cell function) overwhelms the capacity of antioxidant defence and repair system (reviewed by (Monaghan et al., 2009; Pamplona and Costantini, 2011)). This bias towards pro-oxidants inflicts harm to vital cell components (lipids, proteins and DNA), and the accumulation of oxidative damage can play a role in many diseases and ageing-related disorders (Buttemer et al., 2010; Finkel and Holbrook, 2000; Halliwell and Gutteridge, 2007).

Even though glucose is one of the most stable sugars in terms of oxidation, which probably led to the selection of glucose as a primary metabolic compound in the animal kingdom (Bunn and Higgins, 1981), high blood glucose is still a significant risk factor for oxidative stress (Barbieri et al., 2003; Braun and Sweazea, 2008; Ceriello, 2000; Costantini, 2008; Kristal and Yu, 1992). Specifically, high circulating levels of glucose cause an overproduction of electron donors in the tricarboxylic acid cycle, which increases mitochondrial proton gradient and superoxide generation (Du et al., 2000; Nishikawa et al., 2000). Excess superoxide subsequently inhibits glycolysis, activating the four main pathways of glucose over-utilisation known to cause diabetes-associated pathologies (Brownlee, 2001). Two of these pathways, namely the polyol pathway and non-enzymatic glycation further exacerbate oxidative stress by depleting vital intracellular antioxidants – NADPH and reduced glutathione (GSH) – and through the generation of advanced glycation end-products (AGEs; (Brownlee, 2001)). AGEs result from proteins or lipids being cross-linked by dicarbonyls, which arise either as products of direct autoxidation of glucose or from non-enzymatic reactions of glucose-derived metabolites (Barbieri et al., 2003; Brownlee, 2001; Holmes et al., 2001). AGEs can ultimately trigger a cascade of oxidation-mediated damage (Brownlee, 2001; Holmes et al., 2001; Kristal and Yu, 1992). Oxidative stress induced by chronic high blood glucose levels or AGEs predisposes cells and tissues to dysfunctions, leading to varied pathologies in both humans and non-human animals (Braun and Sweazea, 2008; Brownlee, 2001; Ceriello, 1997; Ceriello, 2000; Picard et al., 2014), and may in part be responsible for ageing (Barbieri et al., 2003; Cerami, 1985; Holmes et al., 2001; Kristal and Yu, 1992). It has been shown that hyperglycaemia-induced phenotypes can be ameliorated by antioxidants or overexpression of uncoupling proteins (UCPs), further supporting the involvement of oxidative physiology in diabetes-associated diseases (Ceriello, 1997; Du et al., 2000; Nishikawa et al., 2000). Experimentally induced diabetes in rats increases the level of AGEs, carbonyl stress and oxidative stress (e.g. malondialdehyde), whereas transgenic individuals that overexpress the glyoxalase-I enzyme, which detoxifies AGE precursors, avoid hyperglycaemia-induced oxidative stress (Brouwers et al., 2011).

Despite the considerable implications for health and healthy ageing, surprisingly few studies assessed the functional link between blood glucose and oxidative stress in birds or any other free-living taxa. At the intraspecific level, baseline blood glucose is positively associated with oxidative damage to lipids (malondialdehyde) in house sparrows *Passer domesticus* (Vágási et al., 2020). However, it is still unexplored whether these consequences of high blood glucose levels transcend to the macroevolutionary scale (i.e. whether they can explain variation in oxidative balance among species). The only available evidence comes from the comparison of birds and mammals, showing that birds, on average, evolved lower oxidative stress and longer lifespans, despite their higher levels of blood glucose and AGEs (Baker et al., 2022; Costantini, 2008; Holmes and Ottinger, 2003; Holmes et al., 2001; Satoh, 2021). These observations suggest that blood glucose concentrations are evolutionarily decoupled from oxidative stress levels and thus might not constrain the evolution of long lifespans in birds (see also (Tomášek et al., 2019)). Therefore, it has been proposed that species evolving high blood glucose levels concomitantly evolved adaptations protecting them from (glyc)oxidative damage and resultant cellular dysfunction induced by high blood sugar levels (Braun and Sweazea, 2008; Holmes and Ottinger, 2003; Holmes et al., 2001; Satoh, 2021), which allowed birds in general to evolve a ‘benign hyperglycaemia’ (Baker et al., 2022). Among many adaptations proposed to keep glucose-induced oxidative stress at bay, antioxidants, such as uric acid, glutathione, or the total antioxidant capacity of the plasma, belong to the most commonly considered candidates (Braun and Sweazea, 2008; Brouwers et al., 2011; Ceriello, 1997).

Our understanding of the macroevolutionary association between blood glucose and oxidative stress is hindered by the lack of phylogenetic comparisons across wide ranges of free-living species with diverse life histories. Here, we fill this gap using data from 51 free-living European songbird species sampled during breeding. We collected data on blood glucose levels at physiological baseline and after 30 minutes of standardised stress exposure and computed the magnitude of stress-induced increase in glucose level (termed ‘glucose stress reactivity’ (Vágási et al., 2020)). While the baseline levels reflect the routine energy demands of the species, the stress-induced glucose peaks provide an estimate of the species’ ability to mobilise energy stores depleted during the acute stress response (Romero and Beattie, 2022; Tomášek et al., 2019; Tomášek et al., 2022). The positive correlation between the stress-induced change levels of glucose and corticosterone are often reported ((Jimeno et al., 2018; Vágási et al., 2020); but see exceptions (Deviche et al., 2016; Romero and Beattie, 2022)) and may adaptively be modulated according to life-history strategy (Wingfield and Sapolsky, 2003). Therefore, the macroevolutionary association of both baseline glucose and glucose stress reactivity with oxidative physiology could illuminate the potential costs of evolving high levels of circulating glucose. To assess blood oxidative balance of the species, we measured three non-enzymatic antioxidant parameters (total antioxidant status, uric acid and total glutathione) and the amount of peroxidative damage to lipids (quantified as malondialdehyde levels). Given the hypothesis that high blood glucose levels are detrimental to health partly by causing oxidative stress, we predicted that species that evolved high baseline also evolved higher investment into antioxidant defence or otherwise should experience increased oxidative damage. An alternative hypothesis is that the evolution of blood glucose level and that of oxidative homeostasis is decoupled in birds by adaptations that circumvent the potential adverse consequences of high blood glucose. In this case, we expect no or only weak evidence for the glucose–redox state relationship. There is also a possibility that the stress-induced blood glucose level or glucose stress reactivity may be more important in defining species-specific oxidative physiology than the baseline level. To explore this hypothesis, we examined the relationship between the stress-induced blood glucose parameters and oxidative state variables, despite the fact that the latter are not influenced by capture stress (Vágási et al., 2016).

## MATERIALS AND METHODS

### Field data collection

Samples for blood glucose levels were collected from birds captured in Czech Republic from April to July in 2014 and 2019 (details and data in (Tomášek et al., 2019; Tomášek et al., 2022)), and those for oxidative physiology from birds captured in Romania from April to July between 2011 and 2013, and between 2016 and 2019 (details and data in (Vágási et al., 2016; Vincze et al., 2022)). Depending on the oxidative physiology variable, 50 or 51 species had both glucose and oxidative values (681 individuals with blood glucose and 1021 individuals with oxidative stress data). Samples were collected from adults during the breeding season. Because small-sized songbirds rarely skip breeding (Snow et al., 1998), and thus all individuals in reproductive mode, this sampling period ensures the lowest possible noise in physiological measures. The physiological readiness for breeding was confirmed by the presence of breeding patch in females and cloacal protuberances and sperm production in males (for detailed information, see (Horák et al., 2022; Vágási et al., 2016)). Outside the breeding season, species markedly differ in energy demanding processes, including substantial variation in timing and extent of moulting and/or migratory behaviour (ranging from sedentary to short- and long-distance migration). Therefore, during breeding all small-sized songbird individuals are physiologically in breeding mode, which makes possible to compare physiological traits across species.

### Biochemical parameters

We measured baseline blood glucose concentrations (hereafter G_0_) with FreeStyle Freedom Lite portable glucose meters (Abbott Diabetes Care; linear range: 1.1– 27.8 mM) from samples collected within 3 minutes (baseline glucose levels do not increase within this period following the stress stimulus; (Tomášek et al., 2019)) and stress-induced glucose levels from samples collected after 30 minutes of standardised restraint in cloth bags (hereafter G_30_). G_30_ reflects the ability to mobilise circulating energy in stressful conditions. Species-specific G_0_ and G_30_ values are medians (Tomášek et al., 2019) as they are less affected by outliers than mean values. We computed ‘glucose stress reactivity’ (hereafter ΔG) as a percentage change from G_0_ median to G_30_ median (where G_0_ = 100%) to express the magnitude of change in blood glucose levels when exposed to an acute stress stimulus. The absolute change (i.e. difference between G_0_ and G_30_ medians) is not informative about the magnitude of change in a multispecies dataset, where species largely differ in these values (e.g. change from G_0_ = 5 to G_30_ = 10 mg/dL and from G_0_ = 15 to G_30_ = 20 mg/dL are equal in absolute change [5 mg/dL], while considerably differ in magnitude [100% vs. 33%, respectively]). As expected, mean G_30_ values are significantly higher than G_0_ values in our dataset based on 51 bird species (phylogenetic paired *t*-test, *t* = –5.20, df = 48, *P* < 0.0001) in accordance with a recent result from our extended dataset of 160 bird species (Tomášek et al., 2022).

To assess oxidative state, we measured three non-enzymatic antioxidant markers (total antioxidant status, TAS; uric acid, UA; and total glutathione, tGSH) and a marker of peroxidative damage to membrane lipids (malondialdehyde, MDA). Detailed protocols can be found in (Bókony et al., 2014). Briefly, TAS, UA and MDA were measured from plasma, while tGSH from erythrocytes. TAS level (mM Trolox equivalents) was measured based on a commercial kit (Cayman Chemical, Ann Arbor, MI) according to the method described by (Erel, 2004), with slight modifications as per (Sepp et al., 2010). This assay relies on the ability of non-enzymatic antioxidants (e.g. uric acid, vitamins, sulfhydryl groups of proteins, glutathione) to decolorize the blue-green ABTS^+^ (2,2’-azino-bis(3-ethylbenzothiazoline-6-sulfonate)) to a degree proportional to their concentrations, which can be measured spectrophotometrically at 660 nm (see (Bókony et al., 2014)). UA concentration (mg/dL plasma) was determined spectrophotometrically from 5 μL of plasma by an uricase/peroxidase method (Uric Acid liquicolor kit, Human, Wiesbaden, Germany) (see (Bókony et al., 2014)). MDA is a carbonyl compound that results from the peroxidative degeneration of membrane polyunsaturated fatty acids by reactive oxygen species, and thus it is a widely used marker of oxidative stress (Del Rio et al., 2005). MDA concentration (μg/mL plasma) was determined from 10 μL of plasma by High Performance Liquid Chromatography (HPLC) on a HPLC SUPELCOSIL™ LC-18 column (5 μm particle size; Sigma-Aldrich) with UV detection at 254 nm (Jasco, UV-2075 Plus, Japan) (see (Bókony et al., 2014)). GSH is the most important intracellular, endogenous, non-enzymatic antioxidant (Galván and Alonso-Alvarez, 2008). The concentration of tGSH (nM/mg of erythrocyte pellet) was assayed using a commercial kit (Sigma-Aldrich, St Louis, MO) and according to (Galván and Alonso-Alvarez, 2008) and (Hõrak et al., 2010) with the following modifications (see (Bókony et al., 2014)). Blood samples drawn from small species were usually only enough for one aliquot, and therefore different oxidative state markers were measured from samples from different individuals of the same species resulting in variable sample size for different parameters across species (Vágási et al., 2016).

Due to inter-annual variation in oxidative physiology measures, we computed species-specific values as best linear unbiased estimates (BLUEs; (Vincze et al., 2022)). BLUEs were extracted from linear mixed models (using R function *lmer* from R package *lme4* (Bates et al., 2015)), including each oxidative stress parameter as dependent variables (log (x+1) transformed to ensure model residual normality), as well as species as fixed effect and year of capture as a random grouping variable. There was very strong evidence for positive pairwise associations between species-specific TAS, UA and MDA values (Pearson’s correlation, all *n* = 112, TAS vs. UA: *r* = 0.45, *P* < 0.0001; TAS vs. MDA: *r* = 0.32, *P* = 0.0001; UA vs. MDA: *r* = 0.62, *P* < 0.0001), while there was no evidence for tGSH being associated with the other oxidative parameters. These findings agree with our previous results based on a considerably smaller oxidative physiology dataset (Vágási et al., 2016). We have previously shown that all blood glucose (G_0_, G_30_ and ΔG) and oxidative state variables measured here are suitable for among-species comparative analyses because they fulfil the criterion of being species-specific, as demonstrated by their significant repeatability within species (i.e., conspecifics resemble each other more than members of other species; see details in (Marton et al., 2022; Tomášek et al., 2022)).

### Life-history traits

Most biological traits depend on body size; thus, we used body mass to account for allometry. Species-specific body mass was extracted from (Storchová and Hořák, 2018), but values were verified using alternative sources and data for some species were updated using recent primary literature or our field measurements (see Table S2 in (Vincze et al., 2022)). Because both blood glucose and oxidative physiology coevolve with pace-of-life as expressed by reproductive investment (Tomášek et al., 2019; Vágási et al., 2019), we also computed species-specific annual fecundity (i.e. clutch size × no. clutches per year). Data on clutch size and the number of clutches per year were retrieved from (Vágási et al., 2019).

### Statistical analyses

All statistical analyses were carried out in R 4.0.4 (R Core Team, 2021). To explore the glucose– oxidative physiology relationships across species, we built two sets of phylogenetic generalised least squares (PGLS) models. In the first model set, the BLUEs of the four oxidative physiology parameters were used as response variables in separate models, while body mass and the examined glucose measure (G_0_ or G_30_ or ΔG) were continuous explanatory variables. In the second model set, besides body mass, we also controlled for the potential effect of additional confounding variables. Annual fecundity was included in all models of the second set to rule out the potential confounding effect of pace-of-life. In the models of TAS, UA and MDA, we also included as additional predictors those oxidative parameters (as BLUEs) that correlate with the response variable (e.g. UA and MDA in the TAS model) to rule out the potentially confounding effect of intercorrelating oxidative parameters (see above). All explanatory variables were log-transformed to meet the normality assumption of the model residuals.

Models were constructed using the *gls* function of package *nlme* (Pinheiro et al., 2015). Phylogeny was downloaded from http://birdtree.org (Jetz et al., 2012). We downloaded 1000 random trees using the Hackett backbone (Hackett et al., 2008) and merged them into a rooted, ultrametric consensus tree using the SumTrees Python library (Sukumaran and Holder, 2010). The phylogenetic signal was estimated by maximum likelihood and set to the most appropriate value in each model.

Each model was run with multiple model weights: (1) without weights, (2) weighted by log(sample size +1) and (3) weighted by raw sample size of the dependent variable to control for the sampling effort (see also (Vágási et al., 2019)). Akaike’s Information Criterion (AIC) was used to compare models with different weights. Unweighted models performed best in the case of the UA and MDA, models weighted by log sample size had consistently lower AIC values in case of TAS, while models weighted by sample size provided the best fit in the case of tGSH. Therefore, plots for TAS and tGSH were prepared to show point sizes proportional to either log sample size or sample size, respectively. Fitted lines and associated standard errors on the plots were obtained from the respective full models. Given that physiological and life-history traits often intercorrelate, we verified multicollinearity in the models by computing the variance inflation factors (VIFs) within each full model. All VIFs were below the more conservative < 2 threshold, except the intercorrelating oxidative parameters had higher VIFs, which were below the < 5 threshold (max VIF = 4.01 for MDA in TAS model). Therefore, multicollinearity is unlikely to influence our results. Results are communicated throughout using the language of evidence (i.e. how strong the evidence is for an association) instead of the significance cut-off at *P* ≤ 0.05 as suggested by (Muff et al., 2022).

## Results

In the first model set that controlled for allometry only, there was no or weak evidence for an association between blood glucose and oxidative stress parameters (Table 1, Table S1). There was weak evidence for a negative association of G_30_ and ΔG levels with tGSH, and for a negative association of G_0_ levels with MDA (Table 1, Table S1).

**Table 1.**
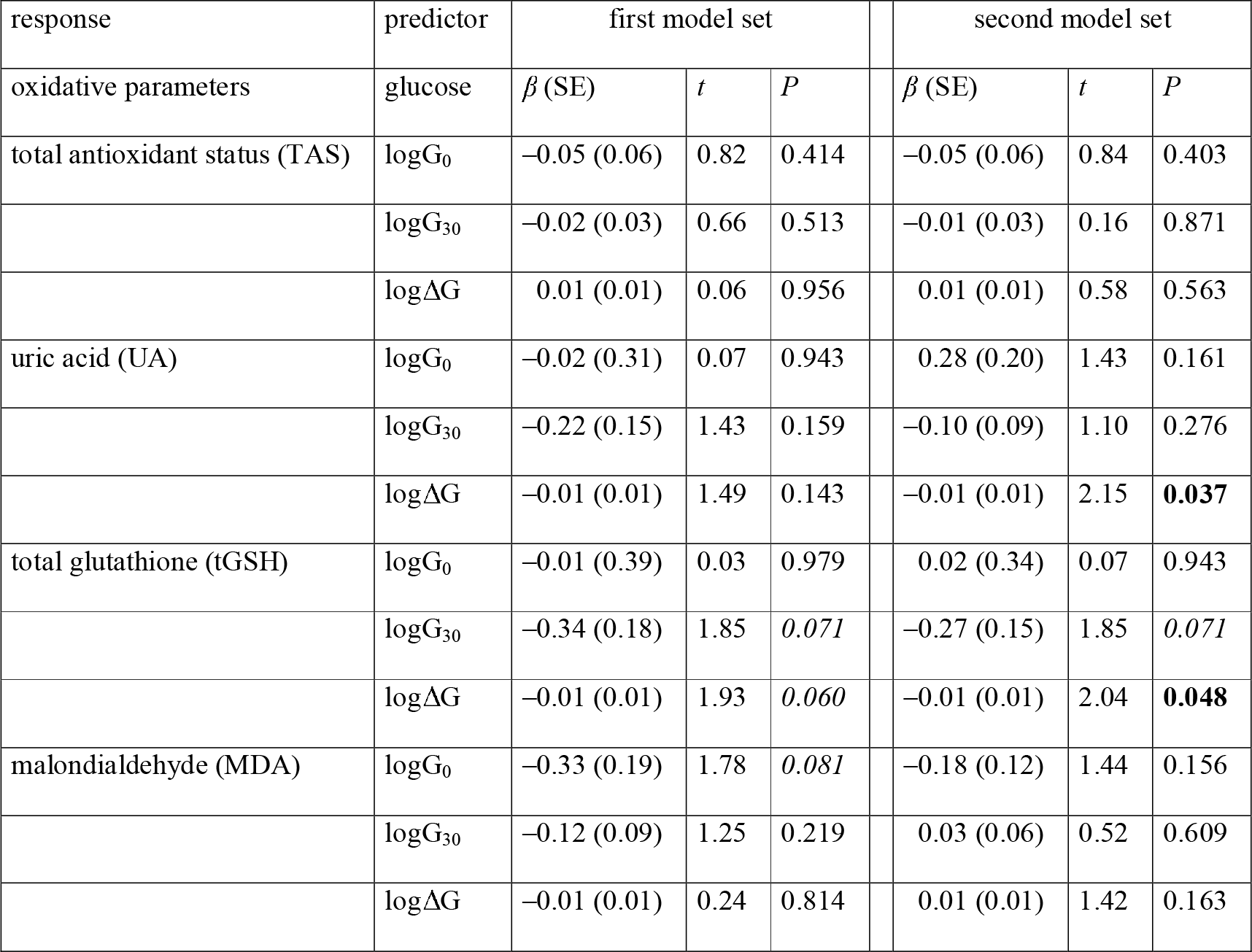
The relationship between oxidative and blood glucose parameters in the first and second sets of phylogenetically controlled models. The first model set analysed the four oxidative parameters in separate models as a function of body mass and glucose as continuous predictors, while the second model set as a function of body mass, fecundity, correlated oxidative parameters and glucose as predictors. Values of oxidative parameters are best linear unbiased predictions (BLUEs, see Methods). Glucose was measured either at baseline (logG_0_), after 30 minutes standardised acute stress exposure (logG_30_) or calculated as % change between G_0_ and G_30_ (logΔG). Full models with all the predictors together with model parameters for the first and second models sets are presented in Table S1 and S2, respectively. *P*-values of associations with moderate evidence (i.e. 0.01 ≤ *P* ≤ 0.05) are marked in bold, while those of weak evidence (i.e. 0.05 ≤ *P* ≤ 0.1) are marked in italics.

The second model set that controlled for allometry, pace-of-life (fecundity) and correlations among oxidative parameters also revealed only a few weak associations between blood glucose and oxidative state. The data revealed only weak evidence for negative associations of ΔG with UA and tGSH levels (Fig. 1b–c), and of G_30_ with tGSH (Table 1, Table S2). There was no evidence of any other associations between glucose and oxidative physiology parameters (Table 1, Table S2, Fig. 1). In this second model set, there was no evidence for an association between G_0_ and MDA (Table 1, Table S2) as compared with the weak association revealed by the first model set (Table 1, Table S1).

**Fig. 1.**
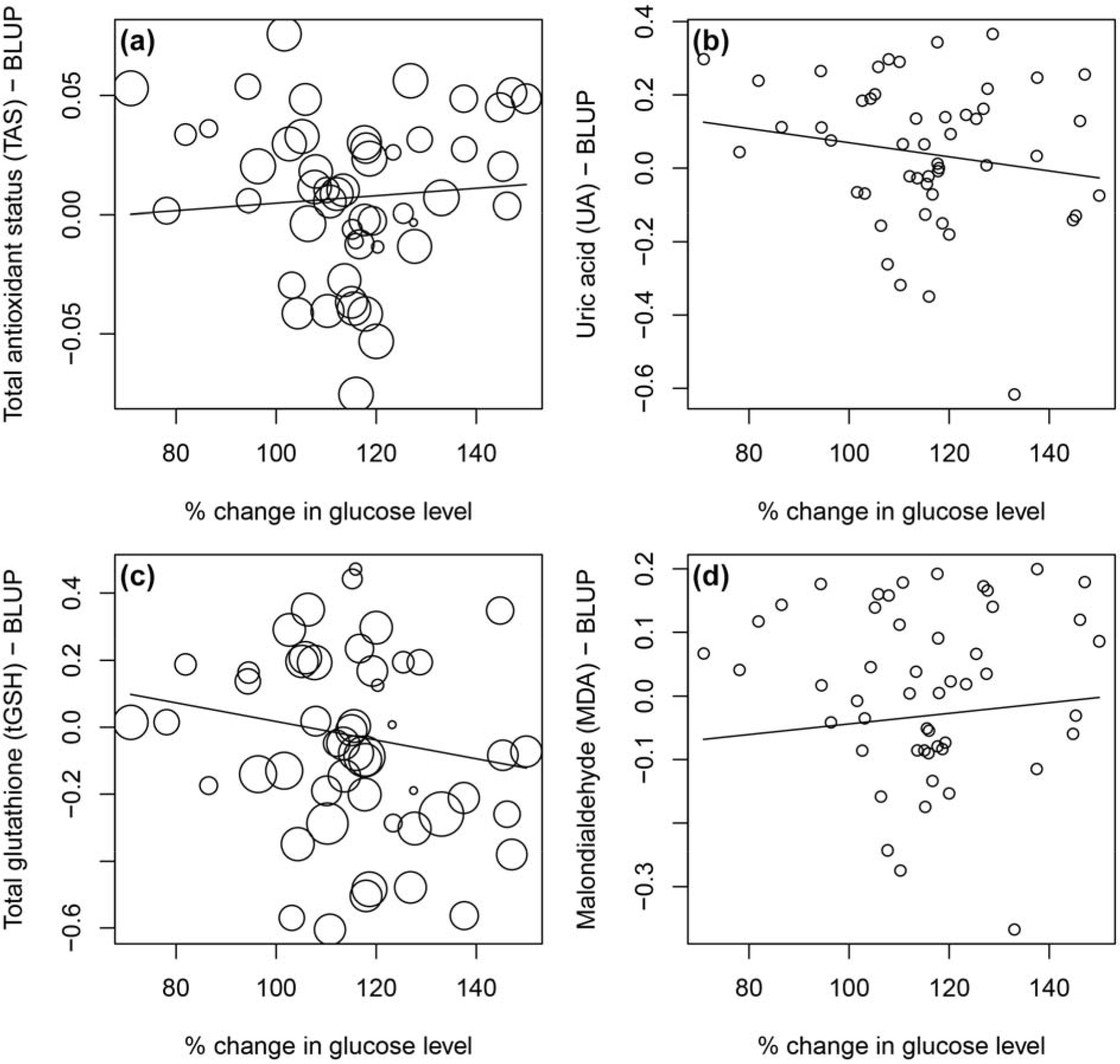
Phylogenetic relationship between glucose stress reactivity (i.e. ΔG, expressed as % change between G_0_ and G_30_, where G_0_ = 100%) and levels of (a) total antioxidant status, (b) uric acid, (c) total glutathione and (d) malondialdehyde. Plots (a) and (c) show point sizes proportional to either log sample size or sample size, respectively (see Methods). Fitted lines were obtained from the respective full models from the second model set (see Table S2).

## Discussion

Our study is the first to examine the oxidative costs (or lack thereof) of high blood glucose levels at the macroevolutionary level. Despite the relatively high blood glucose levels in birds compared with other vertebrates, we found no (or at best only weak) evidence for the macroevolutionary association between blood glucose level and oxidative physiology. The finding that these systems are decoupled supports the hypothesis that birds have evolved adaptations to circumvent the potential oxidative challenges and deleterious consequences imposed by their constitutively high blood glucose levels. Besides such physiological adaptations, life history constraints or confounding effects might also explain this lack of glucose–redox state relationship in birds.

### Physiological adaptations

Birds abound in physiological adaptations that make them excel in physiological homeostasis and longevity. Here, we discuss four such adaptations that might explain the evolutionary dissociation of glucose metabolism and oxidative state. These are higher efficacy of non-enzymatic and structural antioxidant defences (Pamplona and Costantini, 2011), resistance against AGEs (Baker et al., 2022), constitutive activation of nuclear factor erythroid 2-related factor 2 (NRF2; (Castiglione et al., 2020; Satoh, 2021)) and augmented activity of mitochondrial uncoupling proteins (UCPs; (Criscuolo et al., 2005; Stier et al., 2014)).

The high blood glucose levels of birds would be regarded as hyperglycaemia in mammals of similar body size. In humans and mammalian models, hyperglycaemic conditions lead to elevated oxidative stress, which is the central mechanism inducing activation of all the four main pathways known to generate diabetes-associated pathologies (Brownlee, 2001). In mammalian tissues, high activity of one of these pathways, the polyol pathway, depletes NADPH and reduces glutathione (GSH) reserves, which can augment oxidative stress and associated deleterious consequences of diabetes (Brownlee, 2001). In accordance with this mechanism, we found that bird species that evolved higher glucose concentrations or higher glucose stress reactivity during the breeding season also evolved a concomitant lower level of tGSH. Nevertheless, tGSH was only moderately lower in these species and, importantly, not accompanied by higher oxidative damage. This suggests that songbirds with high blood glucose might have evolved higher rates of GSSG-to-GSH regeneration, thereby preventing GSH depletion despite its lower levels and presumably higher glycoxidative load. The investment into the non-enzymatic antioxidant capacity in general was not considerably lower in birds with elevated glucose levels as the relationships between the two physiological systems were mostly absent or at best weak. Besides, birds might have evolved structural adaptations making their tissues more resistant to oxidative damage (Pamplona and Costantini, 2011).

High blood glucose levels can induce oxidative stress by means other than decreased GSH levels. The augmented activity of the polyol pathway in hyperglycaemia can generate AGEs, which in turn can promote oxidative stress by increasing the rate of free radical and highly reactive dialdehyde (e.g. MDA) formation (Brownlee, 2001; Fukami et al., 2005). Remarkably, MDA was not higher but either unrelated or slightly lower in species with higher baseline glucose levels. This result is consistent with recent findings showing that birds have lower lipid peroxidation compared with mammals (Jimenez et al., 2019), despite having higher glucose and AGE levels ((Baker et al., 2022); but see (Anthony-Regnitz et al., 2020)). Birds, therefore, defy the expectation that more AGEs should induce more lipid peroxidation, suggesting that high blood glucose levels and lipid peroxidation (e.g. MDA production) are uncoupled at macroevolutionary scales in birds. This uncoupling could be underpinned by either lower rates of formation of particular AGEs (e.g. serum albumin; (Anthony-Regnitz et al., 2020)), or higher rates of other AGEs formation but also better clearance of these AGEs or a lack of receptors for AGEs (Baker et al., 2022). These physiological and molecular traits are considered avian adaptations that can circumvent the potential glycoxidative toll of elevated AGE levels.

It has been suggested that Neoaves have elevated expression of antioxidant enzymes and hence higher resistance to oxidative challenge due to the constitutive activation of NRF2 (Castiglione et al., 2020; Satoh, 2021). The plausibility of this hypothesis is supported by the observed ability of the overexpressed antioxidant enzyme superoxide dismutase to prevent hyperglycaemia-induced diabetic complications in mammalian tissues (Brownlee, 2001; Du et al., 2000; Nishikawa et al., 2000). Comparative studies for testing these relationships in birds are desirable.

Augmented activity of avian UCPs is another adaptation that can help to quench the oxidative costs of high blood glucose in birds. UCPs are mitochondrial proton channels that can reduce proton gradient across the inner mitochondrial membrane by leaking protons back into the mitochondrial matrix. Given that hyperglycaemia induces oxidative stress by increasing the proton gradient, which leads to excessive superoxide generation, higher activity of UCPs may represent and important protective adaptation. Although broad-scale comparative studies are lacking, the observation of a considerably higher UCP expression in canaries compared to mice provided promising initial evidence for this hypothesis (Slocinska et al., 2010). Moreover, UCPs have been demonstrated to prevent oxidative stress in both birds and mammals (Criscuolo et al., 2005; Stier et al., 2014) and prolong lifespan in mammals (Andrews and Horvath, 2009; Conti et al., 2006).

### Life history constraints and other potential confounding variables

Physiological systems are evolutionarily linked to life history and behavioural traits leading to the emergence of pace-of-life syndrome (Réale et al., 2010; Ricklefs and Wikelski, 2002). The covariations among the traits integrated into the pace-of-life syndrome pose constraints on each other. Fecundity is a key component of pace-of-life. Increased fecundity requires high blood glucose levels to fuel the energetic demands of reproduction, and this might have negative repercussions for oxidative balance. In accordance with these predictions, species with higher fecundity have higher blood glucose concentrations (Tomášek et al., 2019). Additionally, high fecundity may affect oxidative physiology in other ways unrelated to its co-evolution with high blood glucose levels, for example, due to a lower investment into self-maintenance including antioxidant defences (Vágási et al., 2019), which led to our decision to control for its possible confounding effect in the second model set. Nevertheless, our findings suggest that fecundity does not mediate or confound the glucose–redox state relationship as fecundity had little importance in the models.

Species with high baseline glucose levels (i.e. G_0_) showed weakly lower MDA levels indicating a marginally lower rate of lipid peroxidation. This association merits future studies on larger number of bird species or even other taxa. A potential explanation for this weak negative association, other than the candidate adaptations discussed above, could be that there are confounding variables that were not considered in our models, such as a shift in metabolic substrate use in migratory species towards higher utilisation of lipids, rather than glucose, to produce energy during endurance flights (Jenni and Jenni-Eiermann, 1998). In accordance with this hypothesis, migratory passerines were shown to have lower blood glucose levels than sedentary ones during the breeding season (Tomášek et al., 2019; Tomášek et al., 2022), suggesting such a fuel shift might have indeed evolved and became an inherent property of energy metabolism in migratory species even outside the migratory period. Therefore, such a potential shift in fuel utilisation in long-distance migrants could potentially confound the relationship between MDA and G_0_. Nonetheless, adding migration distance as a continuous predictor to the second model set of MDA with G_0_ did not change the relationship between MDA and G_0_ and only resulted in reduced model fit (results not shown).

Besides, UA is a product of protein catabolism and proteins are also utilised as substrate to fuel strenuous non-stop migratory flights. Similarly to MDA models, migration distance did not change the results of UA (results not shown). These findings indicate that potential fuel shift in migratory species probably does not explain our results.

Diet might also confound the results because more AGEs are formed at higher protein and amino acid concentrations (Cerami, 1985). Our previous results support this notion as we showed that carnivorous birds have higher MDA and lower tGSH levels as compared with herbivorous ones (Marton et al., 2022). However, in the present study we could not assess whether the interaction between diet and glucose variables influences markers of oxidative state because our samples only covered songbirds (see (Tomášek et al., 2019)), a group with less variable diet than in the study by (Marton et al., 2022). Nevertheless, diet probably has little effect on our results because a recent analysis on a much larger dataset including 160 temperate and tropical songbird species, showed that only nectarivory/frugivory elevates blood glucose level (Tomášek et al., 2022), but this diet type is missing in our dataset. Additionally, although plasma lipids (triglycerides), which may largely be of dietary origin, might correlate with MDA in some species (Pérez-Rodríguez et al., 2015), this relationship was clearly non-significant in our dataset (see Supplementary Information in (Vágási et al., 2019)). Therefore, we believe that MDA mostly derives from cell membrane lipids rather than from dietary triglycerides in our samples. Finally, a recent review found that diet macro- and micronutrient manipulations generally have no effect on blood glucose levels in birds (Basile et al., 2022).

Besides, any existent causal relationship between blood glucose and oxidative physiology could be masked under some circumstances because the blood glucose dataset and oxidative physiology dataset were collected from different individuals in two different countries. However, the general requirement of any phylogenetic comparative study is that the among-species differences in trait values that evolved over macroevolutionary timescales are larger than the variation at the within-species level (e.g., sex- or population-related variation), ensuring that macroevolutionary relationships among traits can be detected despite the within-species noise. This assumption is supported when the traits of interest show significant within-species repeatability, which we demonstrated for both blood glucose parameters and oxidative physiology parameters in our previous studies (Tomášek et al., 2022; Vágási et al., 2016). Geographical variation, if additive, should not obscure existing macroevolutionary relationships. To mask such relationships, we would need to assume interactions between traits and different environments. While we cannot rule out geographic interactions entirely, they seem unlikely given similar temperate climates and latitudes in both sample locations (only ca 3° latitude difference). Moreover, analysing data from different sources is a standard practice in comparative studies, including those focused on physiological traits. As opposed to many such comparative studies, however, our dataset is based on highly standardised sampling and lab methodology each physiological trait measured in one lab thereby avoiding potentially considerable interlaboratory variation (Fanson et al., 2017).

In conclusion, the glucose-induced oxidative stress hypothesis is mostly unsupported by our results in songbirds, indicating that blood glucose levels and redox state are largely decoupled at the species level. This suggests that bird species with higher blood glucose levels possess adaptations that prevent glucose-induced oxidative stress, which would otherwise constrain the evolution of long lifespans. Such adaptations may also possibly explain the particular case of birds, which evolve considerably longer lifespans compared to mammals, despite having constitutively high blood glucose levels (referred to as benign hyperglycaemia; (Baker et al., 2022)), the latter being evolved to promote the high metabolic rates and hyperathletic capabilities of birds (Satoh, 2021; Sweazea, 2022). Future comparative studies should explore whether components of the insulin/insulin-like growth factor 1 signalling do coevolve, across a broad range of taxa, with oxidative balance, blood glucose and the candidate adaptations potentially preventing their harmful effects (Satoh, 2021). The implied avian adaptations that prevent glucose-induced oxidative stress and pathologies make birds attractive models for diabetes research as they might help discover treatments to ameliorate diabetes-related health problems found in mammals (Hickey et al., 2012; Sweazea, 2022). Despite these adaptations, birds still can develop hyperglycaemia (with values above their already high normal physiological levels) and show signs of diabetes (Sweazea, 2022). We should also emphasize that our results are valid for the blood tissue, while the glucose–redox state relationship might be different in other tissue types, a topic that merits future attention. Future studies should also assess the association between blood glucose levels and oxidative parameters in other vertebrate groups.

## Acknowledgements

We owe a debt of gratitude to Attila Marton, Lőrinc Bărbos, Judit Veres-Szászka, Krisztina Sándor, Attila Fülöp, Lukáš Bobek, Ondřej Kauzál, Jaroslav Cepák, Miroslav Čejka, Kryštof Horák, Martin Janča, Sampath A. Kumar and Pavlína Opatová for their help on the field and Manuela Banciu and Alina Sesarman for biochemical advice. We thank the directorate of the Botanical Garden Cluj-Napoca for permitting bird capturing. Constructive criticism provided by two anonymous reviewers considerably improved the article.

## Author contributions

CIV, PLP, TA and OT conceived the project; CIV, OV, MA, TK, JP, PLP, TA and OT collected the blood samples; LP and JP carried out the biochemical assays; OV analysed the data with input from CIV, PLP, TA and OT; CIV wrote the paper with significant input from OV, ÁZL, PLP, TA and OT. All authors gave final approval for publication and agreed to be accountable for the aspects of work that they conducted.

## Data availability

All data used in this study are archived at Figshare (https://doi.org/10.6084/m9.figshare.24242935.v1).

