## Supplemental tables S1 & S2 for "Songbirds avoid the oxidative stress costs of high blood glucose levels"

### Supporting Information

Tables S1 and S2

**Table S1.** First set of phylogenetically controlled models analysing the four oxidative parameters in separate models (*a*–*d*) with body mass (log BM) and glucose (log Glu) as continuous predictors. Values of oxidative parameters are best linear unbiased estimates (BLUEs, see main text Methods). Different columns show models run with glucose measured either at baseline ($G_0$, column (1)), after 30 minutes standardised acute stress exposure ($G_{30}$, column (2)) or measured as % change between $G_0$ and $G_{30}$ ($\Delta G$, column (3)). Model parameters are provided in the last row of each model ($\lambda$ – Pagel's lambda, AIC – Akaike's Information Criterion, *n* – number of species). *P*-values of associations with moderate or strong evidence (i.e. $0.001 \leq P \leq 0.05$) are marked in bold, while those of weak evidence (i.e. $0.05 \leq P \leq 0.1$) are marked in italics.

| *response* | (1) logGlu = logG$_0$ | | | (2) logGlu = logG$_{30}$ | | | (3) logGlu = log$\Delta$G | | |
|---|---|---|---|---|---|---|---|---|---|
| predictor | $\beta$ (SE) | *t* | *P* | $\beta$ (SE) | *t* | *P* | $\beta$ (SE) | *t* | *P* |
| *(a) total antioxidant status (TAS) model* | | | | | | | | | |
| intercept | 0.14 (0.16) | 0.92 | 0.364 | 0.07 (0.08) | 0.82 | 0.415 | 0.01 (0.04) | 0.33 | 0.745 |
| logBM | –0.01 (0.01) | 0.63 | 0.533 | –0.01 (0.01) | 0.48 | 0.632 | –0.01 (0.01) | 0.30 | 0.766 |
| logGlu | –0.05 (0.06) | 0.82 | 0.414 | –0.02 (0.03) | 0.66 | 0.513 | 0.01 (0.01) | 0.06 | 0.956 |
| model: $\lambda$ = 0.00, AIC = –167.81, *n* = 50 | | | | $\lambda$ = 0.00, AIC = –166.12, *n* = 50 | | | $\lambda$ = 0.00, AIC = –156.53, *n* = 50 | | |
| *(b) uric acid (UA) model* | | | | | | | | | |
| intercept | 0.31 (0.86) | 0.36 | 0.720 | 0.85 (0.45) | 1.89 | 0.065 | 0.50 (0.23) | 2.15 | 0.037 |
| logBM | –0.06 (0.05) | 1.20 | 0.235 | –0.07 (0.05) | 1.47 | 0.148 | –0.06 (0.05) | 1.26 | 0.214 |
| logGlu | –0.02 (0.31) | 0.07 | 0.943 | –0.22 (0.15) | 1.43 | 0.159 | –0.01 (0.01) | 1.49 | 0.143 |
| model: $\lambda$ = 0.59, AIC = –5.71, *n* = 50 | | | | $\lambda$ = 0.57, AIC = –6.29, *n* = 50 | | | $\lambda$ = 0.60, AIC = 2.73, *n* = 50 | | |
| *(c) total glutathione (tGSH) model* | | | | | | | | | |
| intercept | –0.53 (1.07) | 0.49 | 0.625 | 0.38 (0.56) | 0.68 | 0.501 | –0.17 (0.30) | 0.57 | 0.571 |
| logBM | 0.14 (0.07) | 1.94 | *0.058* | 0.13 (0.07) | 1.87 | *0.067* | 0.14 (0.07) | 2.19 | **0.033** |
| logGlu | –0.01 (0.39) | 0.03 | 0.979 | –0.34 (0.18) | 1.85 | *0.071* | –0.01 (0.01) | 1.93 | *0.060* |
| model: $\lambda$ = 0.75, AIC = 16.77, *n* = 51 | | | | $\lambda$ = 0.80, AIC = 14.98, *n* = 51 | | | $\lambda$ = 0.78, AIC = 23.87, *n* = 51 | | |
| *(d) malondialdehyde (MDA) model* | | | | | | | | | |
| intercept | 1.08 (0.51) | 2.12 | 0.039 | 0.51 (0.28) | 1.83 | 0.073 | 0.21 (0.15) | 1.43 | 0.160 |
| logBM | –0.08 (0.03) | 2.32 | **0.025** | –0.06 (0.03) | 2.01 | **0.050** | –0.06 (0.03) | 1.86 | *0.070* |
| logGlu | –0.33 (0.19) | 1.78 | *0.081* | –0.12 (0.09) | 1.25 | 0.219 | –0.01 (0.01) | 0.24 | 0.814 |
| model: $\lambda$ = 0.86, AIC = –52.50, *n* = 50 | | | | $\lambda$ = 0.75, AIC = –49.61, *n* = 50 | | | $\lambda$ = 0.76, AIC = –38.91, *n* = 50 | | |

**Table S2.** Second set of phylogenetically controlled models analysing the four oxidative parameters (TAS – total antioxidant status, UA – uric acid, tGSH – total glutathione, and MDA – malondialdehyde) in separate models (*a*–*d*) with body mass (log BM), fecundity (log Fec), correlating oxidative parameters (TAS and/or UA and/or MDA) and glucose (log Glu) as predictors. Values of oxidative parameters are best linear unbiased estimates (BLUEs, see main text Methods). Different columns show models run with glucose measured either at baseline ($G_0$, column (1)), after 30 minutes standardised acute stress exposure ($G_{30}$, column (2)) or measured as % change between $G_0$ and $G_{30}$ ($\Delta G$, column (3)). Model parameters are provided in the last row of each model ($\lambda$ – Pagel's lambda, AIC – Akaike's Information Criterion, *n* – number of species). *P*-values of associations with moderate or strong evidence (i.e. $0.001 \leq P \leq 0.05$) are marked in bold, while those of weak evidence (i.e. $0.05 \leq P \leq 0.1$) are marked in italics.

| *response* | (1) logGlu = log$G_0$ | | | (2) logGlu = log$G_{30}$ | | | (3) logGlu = log$\Delta G$ | | |
|---|---|---|---|---|---|---|---|---|---|
| predictor | $\beta$ (SE) | *t* | *P* | $\beta$ (SE) | *t* | *P* | $\beta$ (SE) | *t* | *P* |
| *(a) total antioxidant status (TAS) model* | | | | | | | | | |
| intercept | 0.12 (0.15) | 0.81 | 0.422 | 0.01 (0.08) | 0.13 | 0.900 | –0.02 (0.05) | 0.42 | 0.679 |
| logBM | 0.01 (0.01) | 0.06 | 0.951 | 0.01 (0.01) | 0.43 | 0.671 | 0.01 (0.01) | 0.52 | 0.605 |
| logFec | 0.01 (0.01) | 0.11 | 0.913 | –0.01 (0.01) | 0.07 | 0.944 | –0.01 (0.01) | 0.17 | 0.869 |
| UA | 0.05 (0.04) | 1.13 | 0.264 | 0.04 (0.04) | 0.94 | 0.353 | 0.05 (0.04) | 1.11 | 0.272 |
| MDA | 0.04 (0.07) | 0.64 | 0.523 | 0.06 (0.07) | 0.85 | 0.402 | 0.05 (0.07) | 0.69 | 0.494 |
| logGlu | –0.05 (0.06) | 0.84 | 0.403 | –0.01 (0.03) | 0.16 | 0.871 | 0.01 (0.01) | 0.58 | 0.563 |
| model: $\lambda$ = 0.00, AIC = –160.71, *n* = 50 | | | | $\lambda$ = 0.00, AIC = –158.48, *n* = 50 | | | $\lambda$ = 0.00, AIC = –149.66, *n* = 50 | | |
| *(b) uric acid (UA) model* | | | | | | | | | |
| intercept | –0.86 (0.52) | 1.65 | 0.107 | 0.14 (0.28) | 0.50 | 0.617 | 0.09 (0.16) | 0.58 | 0.564 |
| logBM | 0.04 (0.03) | 1.35 | 0.183 | 0.01 (0.03) | 0.48 | 0.630 | 0.02 (0.03) | 0.65 | 0.522 |
| logFec | 0.03 (0.04) | 0.79 | 0.433 | 0.06 (0.04) | 1.41 | 0.166 | 0.05 (0.04) | 1.38 | 0.174 |
| TAS | 0.61 (0.54) | 1.13 | 0.264 | 0.51 (0.54) | 0.94 | 0.353 | 0.58 (0.52) | 1.11 | 0.272 |
| MDA | 1.27 (0.15) | 8.64 | < **0.001** | 1.25 (0.15) | 8.38 | < **0.001** | 1.24 (0.14) | 8.67 | < **0.001** |
| logGlu | 0.28 (0.20) | 1.43 | 0.161 | –0.10 (0.09) | 1.10 | 0.276 | –0.01 (0.01) | 2.15 | **0.037** |
| model: $\lambda$ = 0.00, AIC = –50.37, *n* = 50 | | | | $\lambda$ = 0.00, AIC = –48.03, *n* = 50 | | | $\lambda$ = 0.00, AIC = –42.05, *n* = 50 | | |
| *(c) total glutathione (tGSH) model* | | | | | | | | | |
| intercept | –0.47 (0.86) | 0.55 | 0.588 | 0.28 (0.43) | 0.64 | 0.523 | –0.08 (0.28) | 0.29 | 0.773 |
| logBM | 0.12 (0.05) | 2.17 | **0.035** | 0.11 (0.05) | 2.10 | **0.041** | 0.12 (0.05) | 2.30 | **0.026** |
| logFec | 0.01 (0.08) | 0.14 | 0.893 | 0.03 (0.07) | 0.44 | 0.662 | 0.01 (0.07) | 0.04 | 0.968 |
| logGlu | 0.02 (0.34) | 0.07 | 0.943 | –0.27 (0.15) | 1.85 | *0.071* | –0.01 (0.01) | 2.04 | **0.048** |
| model: $\lambda$ = 0.82, AIC = –1.68, *n* = 51 | | | | $\lambda$ = 0.89, AIC = –3.14, *n* = 51 | | | $\lambda$ = 0.89, AIC = 5.54, *n* = 51 | | |
| *(d) malondialdehyde (MDA) model* | | | | | | | | | |
| intercept | 0.65 (0.32) | 2.02 | 0.050 | 0.12 (0.18) | 0.71 | 0.482 | 0.10 (0.10) | 1.02 | 0.311 |
| logBM | –0.04 (0.02) | 2.52 | **0.016** | –0.03 (0.02) | 2.05 | **0.046** | –0.03 (0.02) | 2.15 | **0.037** |

| | | | | | | | | | | | |
|---|---|---|---|---|---|---|---|---|---|---|---|
| logFec | –0.04 (0.03) | 1.61 | 0.114 | | –0.06 (0.02) | 2.37 | **0.022** | | –0.06 (0.02) | 2.41 | **0.020** |
| TAS | 0.22 (0.33) | 0.66 | 0.516 | | 0.29 (0.34) | 0.85 | 0.402 | | 0.23 (0.34) | 0.69 | 0.494 |
| UA | 0.49 (0.06) | 8.49 | < **0.001** | | 0.49 (0.06) | 8.38 | < **0.001** | | 0.51 (0.06) | 8.67 | < **0.001** |
| logGlu | –0.18 (0.12) | 1.44 | 0.156 | | 0.03 (0.06) | 0.52 | 0.609 | | 0.01 (0.01) | 1.42 | 0.163 |
| model: $\lambda$ = 0.13, AIC = –89.10, *n* = 50 | | | | | $\lambda$ = 0.00, AIC = –87.98, *n* = 50 | | | | $\lambda$ = 0.00, AIC = –80.53, *n* = 50 | | |
